# The association between weight at birth and breast cancer risk revisited using Mendelian randomisation

**DOI:** 10.1101/464115

**Authors:** Siddhartha P. Kar, Irene L. Andrulis, Hermann Brenner, Stephen Burgess, Jenny Chang-Claude, Daniel Considine, Thilo Dörk, D. Gareth R. Evans, Manuela Gago-Domínguez, Graham G. Giles, Mikael Hartman, Dezheng Huo, Rudolf Kaaks, Jingmei Li, Artitaya Lophatananon, Sara Margolin, Roger L. Milne, Kenneth R. Muir, Håkan Olsson, Kevin Punie, Paolo Radice, Jacques Simard, Rulla M. Tamimi, Els Van Nieuwenhuysen, Camilla Wendt, Wei Zheng, Paul D. P. Pharoah

**Affiliations:** Department of Public Health and Primary Care, University of Cambridge, Cambridge, UK; Fred A. Litwin Center for Cancer Genetics, Lunenfeld-Tanenbaum Research Institute of Mount Sinai Hospital, Toronto, ON, Canada; Department of Molecular Genetics, University of Toronto, Toronto, ON, Canada; Division of Clinical Epidemiology and Ageing Research, German Cancer Research Center (DKFZ), Heidelberg, Germany; German Cancer Consortium (DKTK), German Cancer Research Center (DKFZ), Heidelberg, Germany; Division of Preventive Oncology, German Cancer Research Center (DKFZ) and National Center for Tumor Diseases (NCT), Heidelberg, Germany; MRC Biostatistics Unit, University of Cambridge, Cambridge, UK; Division of Cancer Epidemiology, German Cancer Research Center (DKFZ), Heidelberg, Germany; University Medical Center Hamburg-Eppendorf, Cancer Epidemiology, University Cancer Center Hamburg (UCCH), Hamburg, Germany; Gynaecology Research Unit, Hannover Medical School, Hannover, Germany; Division of Evolution and Genomic Sciences, School of Biological Sciences, Faculty of Biology, Medicine, and Health, University of Manchester, Manchester Academic Health Science Centre, Manchester, UK; Genomic Medicine Group, Galician Foundation of Genomic Medicine, Complejo Hospitalario Universitario de Santiago, Servicio Galego de Saude (SERGAS), Instituto de Investigación Sanitaria de Santiago de Compostela (IDIS), Santiago De Compostela, Spain; Moores Cancer Center, University of California San Diego, La Jolla, CA, USA; Cancer Epidemiology and Intelligence Division, Cancer Council Victoria, Melbourne, VIC, Australia; Center for Epidemiology and Biostatistics, Melbourne School of Population and Global Health, University of Melbourne, Parkville, VIC, Australia; Saw Swee Hock School of Public Health, National University of Singapore, Singapore; Department of Surgery, National University of Singapore, Singapore; Department of Public Health Sciences, University of Chicago, Chicago, IL, USA; Human Genetics, Genome Institute of Singapore, Singapore, Singapore; Division of Population Health, Health Services Research and Primary Care, School of Health Sciences, Faculty of Biology, Medicine and Health, University of Manchester, Manchester, UK; Department of Oncology, Södersjukhuset and Department of Clinical Science and Education, Södersjukhuset, Karolinska Institutet, Stockholm, Sweden; Department of Cancer Epidemiology, Clinical Sciences, Lund University, Lund, Sweden; Department of Oncology, Leuven Multidisciplinary Breast Cancer, University Hospital Leuven, KU Leuven, Leuven, Belgium; Unit of Molecular Bases of Genetic Risk and Genetic Testing, Department of Research, Fondazione IRCCS (Istituto Di Ricovero e Cura a Carattere Scientifico) Istituto Nazionale dei Tumori (INT), Milan, Italy; Genomics Center, Centre Hospitalier Universitaire de Québec - Université Laval Research Center, Québec City, QC, Canada; Department of Epidemiology, Harvard T.H. Chan School of Public Health, Boston, MA, USA; Channing Division of Network Medicine, Department of Medicine, Brigham and Women’s Hospital and Harvard Medical School, Boston, MA, USA; Division of Epidemiology, Department of Medicine, Vanderbilt Epidemiology Center, Vanderbilt-Ingram Cancer Center, Vanderbilt University School of Medicine, Nashville, TN, USA; Department of Oncology, University of Cambridge, Cambridge, UK

**Keywords:** birth weight, breast cancer, Mendelian randomisation

## Abstract

Observational studies suggest that higher birth weight (BW) is associated with increased risk of breast cancer in adult life. We conducted a two-sample Mendelian randomisation (MR) study to assess whether this association is causal. Sixty independent single nucleotide polymorphisms (SNPs) known to be associated at *P* < 5 × 10^-8^ with BW were used to construct (1) a 41-SNP instrumental variable (IV) for univariable MR after removing SNPs with pleiotropic associations with other breast cancer risk factors and (2) a 49-SNP IV for multivariable MR after filtering SNPs for data availability. BW predicted by the 41-SNP IV was not associated with overall breast cancer risk in inverse-variance weighted (IVW) univariable MR analysis of genetic association data from 122,977 breast cancer cases and 105,974 controls (odds ratio = 0.86 per 500 g higher BW; 95% confidence interval: 0.73—1.01). Sensitivity analyses using four alternative methods and three alternative IVs, including an IV with 59 of the 60 BW-associated SNPs, yielded similar results. Multivariable MR adjusting for the effects of the 49-SNP IV on birth length, adult height, adult body mass index, age at menarche, and age at menopause using IVW and MR-Egger methods provided estimates consistent with univariable analyses. Results were also similar when all analyses were repeated after restricting to estrogen receptor-positive or -negative breast cancer cases. Point estimates of the odds ratios from most analyses performed indicated an inverse relationship between genetically-predicted BW and breast cancer. Thus, there is little evidence from MR to suggest that the previously observed association between higher BW and increased risk of breast cancer in adult life is causal.

## Background

The hypothesis that the risk of developing breast cancer in adulthood may be increased by factors that first act *in utero* – in particular by fetal exposure to higher levels of maternal estrogen – was first proposed in 1990 [1]. Since then, several observational studies have examined this hypothesis by using birth weight as an index of the effects of intrauterine hormones on fetal growth and of the extent of the fetal mammary stem cell pool from which breast tumours may eventually arise [2—6]]. When considered overall, these studies have suggested that higher weight at birth may be associated with an increased susceptibility to breast cancer in later life [7—20]], but a few studies have failed to demonstrate this association [21—28]]. While most studies have adjusted for the effects of recognised breast cancer risk factors measured at a specific time point in their samples, this does not rule out the possibility of residual effects of these factors, which may act at different points over the course of life, driving the observed association between birth weight and breast cancer. It has not been possible to determine whether this association is causal and to dissect whether it is birth weight *per se* or an external factor influencing fetal growth and development that might underpin a possible association with future breast cancer risk.

Mendelian randomisation (MR) is a form of instrumental variable (IV) analysis that uses genetic instruments or single nucleotide polymorphisms (SNPs) associated with an exposure of interest to infer a causal relationship (or lack thereof) between this exposure and an outcome. Since SNPs are randomly allocated at conception, an MR study may be thought of as being analogous to a randomised controlled trial of the effects of the exposure on the outcome, making such a study less susceptible to classical confounding. However, MR is a valid tool for causal inference only if three assumptions hold: (1) SNPs used as part of the IV are associated with the exposure, (2) these SNPs are not associated with known or unknown confounders for the outcome, and (3) given these confounders, the SNPs affect the outcome only through the exposure of interest and not via other pathways. Here, we report the result of an MR analysis that aimed to investigate the relationship between birth weight as the exposure and susceptibility to breast cancer as the outcome. A genome-wide association meta-analysis of over 150,000 individuals has previously identified 60 independent SNP loci that are associated with birth weight (BW) and genetic association data for 59 of these loci are available from a separate meta-analysis of breast cancer susceptibility that included over 225,000 women [29,30]. We used these data to perform an MR study with a two-sample design (Figure 1) wherein the genetic associations with outcome and exposure are estimated in independent studies. It is challenging to evaluate empirically whether the assumptions required for causal inference with MR are met and therefore, we also report the results of several additional analyses performed to assess the robustness of our MR result.

**Figure 1:**
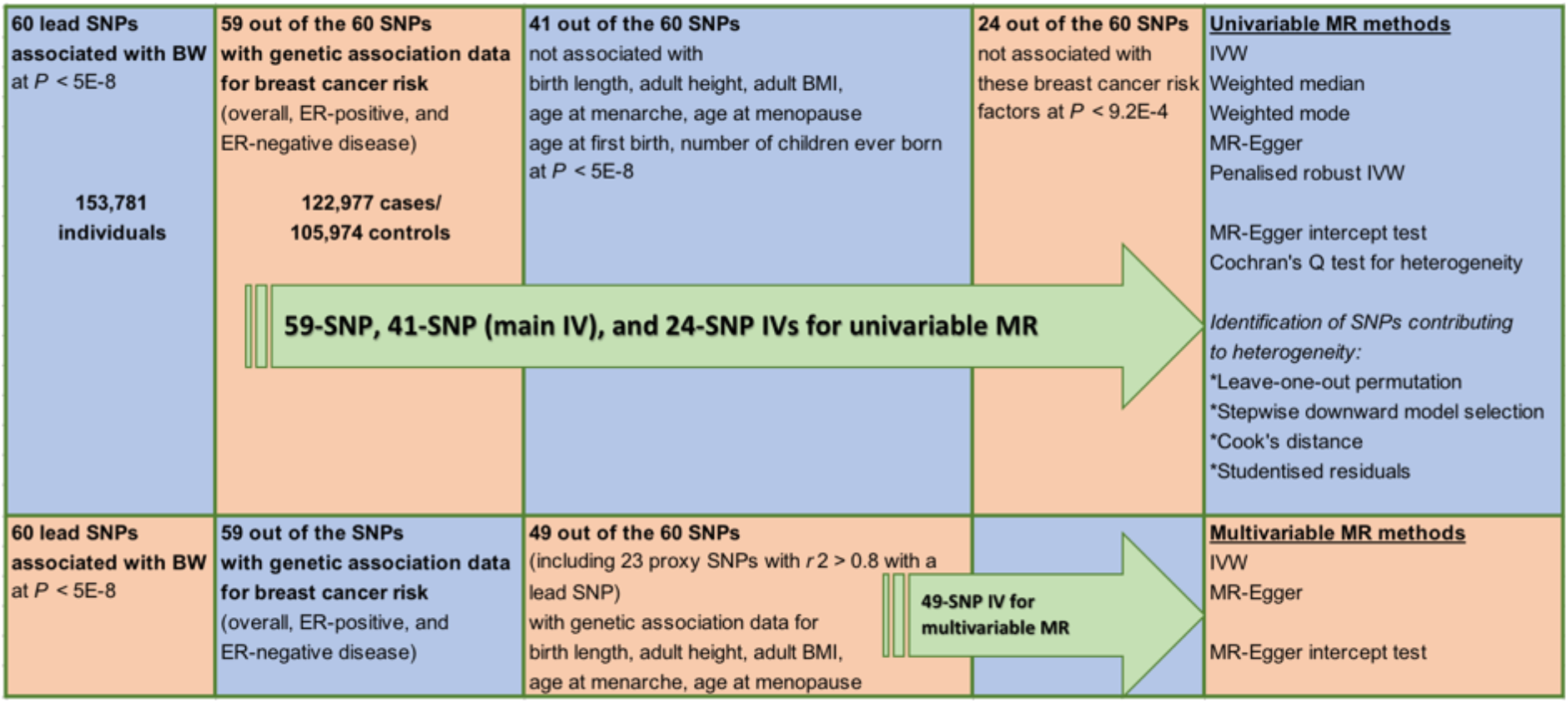
Schematic overview of study design. Abbreviations: BW = birth weight; BMI = body mass index; MR = Mendelian randomisation; IVW = inverse-variance weighted method; SNP = single nucleotide polymorphism; ER = estrogen receptor; IV = instrumental variable.

## Methods

### Birth weight (exposure) data

Genome-wide significant (*P* < 5 × 10^-8^) associations for BW at 60 independent loci were previously reported by Horikoshi et al [29]. The lead SNPs at 49 loci were identified at *P* < 5 × 10^-8^ in the European ancestry component (n = 143,677) of the study by Horikoshi et al. The lead SNPs at the remaining loci had *P*-values ranging from 5.1 × 10^-8^ to 4.4 × 10^-7^ in the European ancestry meta-analysis and were identified at *P* < 5 × 10^-8^ in the trans-ancestry component of the same study, which combined the European results with results from 10,104 individuals of diverse ancestry [29]. Summary results from the European-only meta-analysis for all 60 SNPs were obtained from the Early Growth Genetics (EGG) Consortium (Supplementary Tables 1 and 2). BW in the EGG Consortium studies was collected from heterogeneous sources (from measurements made at birth by medical doctors, birth records and medical registers, maternal interviews, and adult self-report) [29].

### Breast cancer (outcome) data

Summary results from genome-wide association meta-analyses for breast cancer susceptibility in women of European ancestry by Michailidou et al [30] were downloaded from the Breast Cancer Association Consortium (BCAC) website (Supplementary Table 1). Results were available for 59 of the 60 BW-associated lead SNPs (not available for rs139975827) for three breast cancer susceptibility phenotypes (Supplementary Table 2): overall breast cancer risk (122,977 cases/105,974 controls), estrogen receptor (ER)-positive breast cancer risk (69,501 cases/105,974 controls), and ER-negative breast cancer risk (21,468 cases/105,974 controls). No proxy SNPs correlated with rs139975827 with *r*^2^ > 0.8 could be identified. Fifty-eight SNPs were either genotyped or imputed with quality score > 0.8 in the OncoArray project, which was the largest single component of the BCAC meta-analyses [30]. One SNP, rs138715366, was imputed with quality score = 0.62 and also retained for subsequent analyses.

### Birth length and other breast cancer-risk factor data

Summary results from genome-wide association meta-analyses for birth length (n = 28,459) [31], adult height (n = 253,288) [32], adult body mass index (BMI) (in women; n = 171,977) [33], age at menarche (n = 132,989) [34], age at menopause (n = 69,360) [35], age at first birth (in women; n = 189,656) [36], and number of children ever born (in women; n = 225,230) [36] were obtained from four genetic consortia. The consortia and corresponding websites used for data download are listed in Supplementary Table 1. The age at menarche sample used here excluded women genotyped in the Collaborative Oncological Gene-environment Study (COGS) who were included in the breast cancer meta-analyses [30,34]. All summary statistics obtained were from analyses limited to individuals of European ancestry. These statistics included standardised regression (beta) coefficients and *P*-values from genetic association analysis in each data set, with the exceptions of age at first birth and number of children ever born, where only *P*-values were available.

### Data harmonisation

The genetic association data for BW and breast cancer susceptibility were based on imputation using the 1000 Genomes reference panel, while all other data sets had been imputed using HapMap. Out of the 59 BW-associated lead SNPs that were also present in the breast cancer data set, summary statistics were available for 26 SNPs across all data sets (Supplementary Table 3). For an additional 26 SNPs, proxy SNPs in linkage disequilibrium with the BW-associated lead SNP (*r*^2^ > 0.6 in European samples) were previously identified by Horikoshi et al [29]. The same proxies were used here because they were available across all data sets (Supplementary Table 3). For a further two SNPs (rs10402712 and rs7402982), proxies with *r*^2^ > 0.6 in European samples (rs11667352 and rs2017500, respectively) were identified in the current study using the SNP Annotation and Proxy (SNAP) Search tool version 2.2 (http://archive.broadinstitute.org/mpg/snap; [37]). No suitable proxy SNP was identified for four BW-associated lead SNPs (rs61830764, rs138715366, rs72851023, and rs144843919). A fifth SNP (rs11096402) is an X-chromosome variant mapped only in the BW and breast cancer data sets.

In summary, breast cancer data were available for 59 of the 60 BW-associated lead SNPs. When considering these two data sets together with the seven additional “risk factor” data sets, 54 BW loci had results listed across all data sets [26 represented by BW-associated lead SNPs and 28 by correlated (*r*^2^ > 0.6) proxy SNPs]. The signs of the beta coefficients for each SNP across all summary data sets were aligned to the BW-increasing allele (except for age at first birth and number of children ever born, where only *P*-values were available).

### Univariable Mendelian randomisation

Of the 54 BW loci represented across all data sets, 13 demonstrated association with at least one of the seven breast cancer risk factors at *P* < 5 × 10^-8^ (Supplementary Table 3). The BW-associated lead SNP at the 41 loci that did not show genome-wide significant pleiotropic associations were used to construct the main 41-SNP IV for univariable MR analyses (these SNPs are highlighted in Supplementary Table 2). This was done to ensure that the univariable MR analyses were, as far as possible, evaluating the association between BW and adult breast cancer risk independent of the effects of SNPs that were also associated with other potential later-life breast cancer risk factors such as adult height. MR was first performed using the inverse-variance weighted (IVW) method wherein SNP associations with the outcome (breast cancer) were regressed on SNP associations with the exposure (BW) using a linear model weighted by the inverse of the variance of SNP associations with the outcome [38]. This approach to MR analysis as applied to summary genetic association statistics is equivalent to standard two-stage least squares regression MR using individual-level genetic data [38]. The standard error of the IVW effect was estimated by a multiplicative random-effects model.

The IVW method assumes that the IV as a whole satisfies the MR assumptions or that all SNPs in the IV constitute valid instruments. Because the validity of this assumption is difficult to test empirically, four additional MR methods were also used as sensitivity analyses to assess the robustness of the result from the primary analysis under alternative assumptions: weighted median function [39], weighted mode function [40], MR-Egger regression [41], and penalised robust IVW regression [42]. The weighted median function provides a valid result under the assumption that over 50% of the weight in the IV model comes from SNPs that satisfy the MR assumptions. The weighted mode function is valid if the largest number of SNPs with similar individual Wald ratios [beta(outcome)/beta(exposure)] are valid instruments even if other SNPs in the IV do not meet the requirements for causal inference using MR. The weighted effect in these functions is derived from the inverse of the standard error of the Wald ratio for each SNP. Penalised robust IVW uses robust regression in place of standard linear regression-based IVW, with a penalty on the contribution to the analysis of SNPs with outlying Wald ratios. MR-Egger regression allows for horizontal pleiotropic effects, i.e., the association between SNPs in the IV and other traits that may affect the outcome via pathways independent of the exposure. It assumes that such pleiotropy is not correlated with SNP-exposure associations and under this assumption, the MR-Egger result is valid even if all SNPs in the IV are invalid instruments for MR due to their pleiotropic associations. Further, the MR-Egger regression intercept provides an estimate of the average pleiotropic effect over all SNPs in the IV and if this differs from zero, it indicates the presence of horizontal pleiotropy [41]. The null hypothesis that this intercept did not differ significantly from zero was also tested. The R package *MendelianRandomization* (v0.2.2) was used for the penalised robust IVW MR analysis and *TwoSampleMR* for all other MR methods [42,43].

Pleiotropy was also assessed by calculating Cochran’s Q as a measure of between-instrument heterogeneity using the Wald ratios for each SNP in the 41-SNP IV [44]. To identify specific SNPs in this IV that contributed to observed heterogeneity and were potentially pleiotropic, four steps were taken. First, SNPs were removed iteratively from the 41-SNP IV until Cochran’s Q test was no longer significant at the *P* < 0.05 level using the stepwise downward “model selection” procedure implemented in the R package *gtx* (v0.0.8). Second, Cook’s distance (with a threshold of 4/number of SNPs in the IV) was used to identify SNPs in the 41-SNP IV with a disproportionate influence on the primary IVW model [45,46]. Third, leave-one-out permutation analysis was performed, i.e., each SNP was sequentially removed from the 41-SNP IV and the primary IVW MR analysis repeated. Fourth, studentised residuals (with a threshold of ±2) were used to identify outlier SNPs in the IVW model [46]. The effect of pleiotropy was further investigated by applying the standard IVW approach to a 24-SNP IV created by removing all BW loci associated with at least one of the seven breast cancer risk factors at *P* < 9.2 × 10^-4^ (after Bonferroni correction for examining pleiotropic associations at 54 BW loci). As with the main 41-SNP IV, all 24 SNPs were BW-associated lead SNPs (the SNPs are highlighted in Supplementary Table 2). Finally, we note that we did also test using the IVW method the full 59-SNP IV (i.e., all SNPs listed in Supplementary Table 2), which constituted all known BW-associated lead SNPs with matched breast cancer data available regardless of their pleiotropic associations.

The proportions of variance of BW explained by the 41-SNP and 59-SNP IVs were estimated using the “steiger.R” function (https://rdrr.io/github/MRCIEU/TwoSampleMR/src/R/steiger.R) and *a priori* power to detect an association at a significance level of 0.05 was calculated using an online tool (https://sb452.shinyapps.io/power) [47]. All statistical tests were 2-sided and *P* < 0.05 was considered statistically significant unless otherwise specified. Each analysis was repeated for the three outcomes: overall breast cancer risk, ER-positive breast cancer risk, and ER-negative breast cancer risk.

### Multivariable Mendelian randomisation

Multivariable MR (MVMR) represents an alternative strategy for conducting an MR analysis using an IV that contains SNPs which in addition to their established association with the exposure of interest are also associated with other known risk factors for the outcome [48,49]. MVMR allows for the inclusion of such SNPs in the IV by adjusting for their associations with these risk factors. MVMR analysis was performed by regressing SNP associations with breast cancer on SNP associations with BW, birth length, adult height, adult BMI, age at menarche, and age at menopause in a single regression model. This model thus helped estimate the effect of BW on breast cancer independent of the effects of these other factors on breast cancer [48,49]. Like its univariable counterpart, an inverse-variance weighted linear regression model with multiplicative random effects was used for MVMR. The IV used for MVMR contained 49 SNPs (Supplementary Table 4). As described under “Data harmonisation”, 26 SNPs were BW-associated lead SNPs and 23 were proxy SNPs strongly correlated (*r*^2^ > 0.8) with the BW-associated lead SNPs at their respective BW loci (five additional proxy SNPs with *r*^2^ > 0.6 but *r*^2^ < 0.8 were omitted from the MVMR IV; Supplementary Tables 3 and 4). Since beta coefficients were not available for SNP associations with age at first birth and number of children ever born, these variables were not included in the MVMR model. However, none of the 49 SNPs were associated at *P* < 5 × 10^-8^ with these two variables (Supplementary Tables 3 and 4). Multivariable extensions of MR-Egger regression and the MR-Egger intercept test were also applied to this 49-SNP IV [50]. Each analysis was repeated for the three outcomes: overall breast cancer risk, ER-positive breast cancer risk, and ER-negative breast cancer risk.

### Statistical power

The 41-SNP IV explained 1.3% of the variance of BW and had over 80% power to detect 11% increase (or decrease) in overall breast cancer risk in the univariable MR analysis (odds ratios (ORs) of 0.89 or 1.11 per 1-standard deviation (SD; 500 g) higher BW). The 59-SNP IV explained 2% of the variance of BW and had over 80% power to detect 9% increase (or decrease) in overall breast cancer risk in the univariable MR analysis (ORs of 0.91 or 1.09 per 1-SD higher BW). For comparison, the largest observational investigation of the association between BW and breast cancer risk [7], a pooled analysis of individual participant data from 32 studies (22,058 breast cancer cases and 604,854 non-cases) identified 6% increase in breast cancer risk or a pooled relative risk per 1-SD (500 g) higher BW of 1.06 (95% confidence interval (CI): 1.02—1.09). The 41-SNP and 59-SNP genetic instruments had 35% and 50% power, respectively, to detect an OR of 1.06. Power calculations for the corresponding univariable (IVW) MR analyses specific to ER-positive and ER-negative breast cancer risks are provided in Supplementary Table 5. We also confirmed the power of the 41-SNP IV to detect an association between lower BW and increased risk of type 2 diabetes (T2D) in later life using summary genome-wide association meta-analysis statistics from 26,676 T2D cases and 132,532 controls (Supplementary Table 1; IVW OR = 1.93; 95% CI: 1.20—3.08; *P* = 0.006) [51]. This association has been previously identified in two MR studies [52,53].

## Results

### Associations between individual SNPs and breast cancer

Of the 59 BW-associated lead SNPs, 18 were associated with overall breast cancer risk at *P* < 0.05, six after Bonferroni correction for testing 59 SNPs (*P* < 8 × 10^-4^), and two at *P* < 5 × 10^-8^ (Supplementary Table 2). Of the 18 SNPs associated with overall breast cancer risk at *P* < 0.05, the direction of association with BW was inverse for 13 SNPs (i.e., the allele that increased BW was protective for breast cancer). Notably, this direction of association is contrary to that identified by observational studies which suggest that higher BW is associated with increased breast cancer risk in later life. The 13 SNPs included both genome-wide significant overall breast cancer risk SNPs, rs2229742 (encoding missense mutation R448G in *NRIP1*) and rs1101081 (intronic SNP in *ESR1*). The 18 SNPs represented a six-fold enrichment over the number of associations with breast cancer at *P* < 0.05 that were expected by chance alone. Associations between individual SNPs and ER-positive and ER-negative breast cancer are also provided in Supplementary Table 2.

### Univariable Mendelian randomisation

MR analysis of the 41-SNP IV using the IVW method yielded an OR of 0.86 (95% CI: 0.73—1.01; *P* = 0.06) for overall breast cancer per 1-SD (500 g) higher BW (Figure 2 (a)). These estimates were consistent in direction with the results of the weighted median, weighted mode, MR-Egger, and penalised robust IVW sensitivity analyses (Figure 2 (a) and Supplementary Figure 1 (a)). The MR-Egger intercept test (*P* = 0.26) suggested absence of strong directional horizontal pleiotropy (i.e., effects of the 41-SNP IV on breast cancer via a pathway that arises proximal to or upstream of BW).

**Figure 2:**
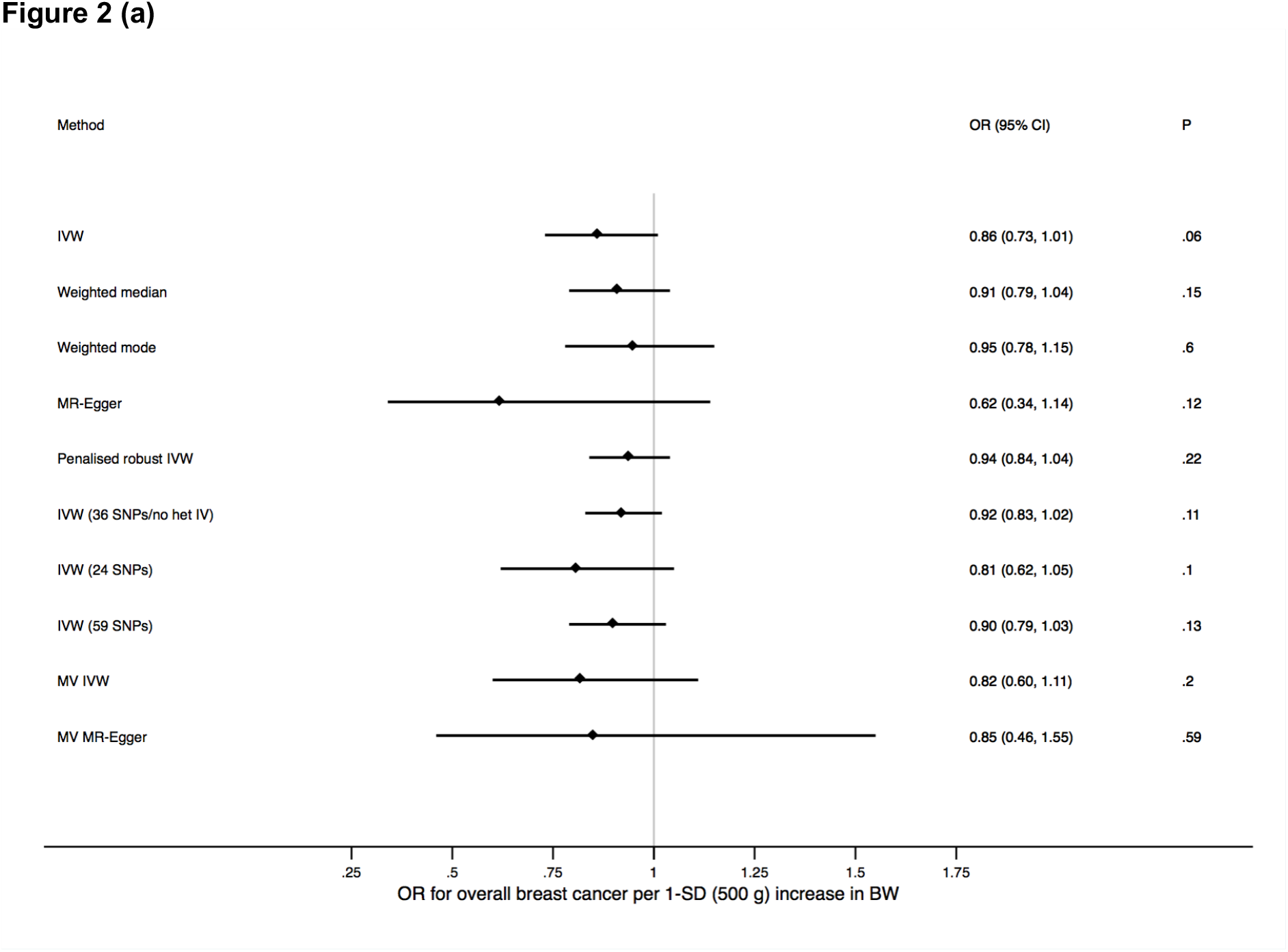
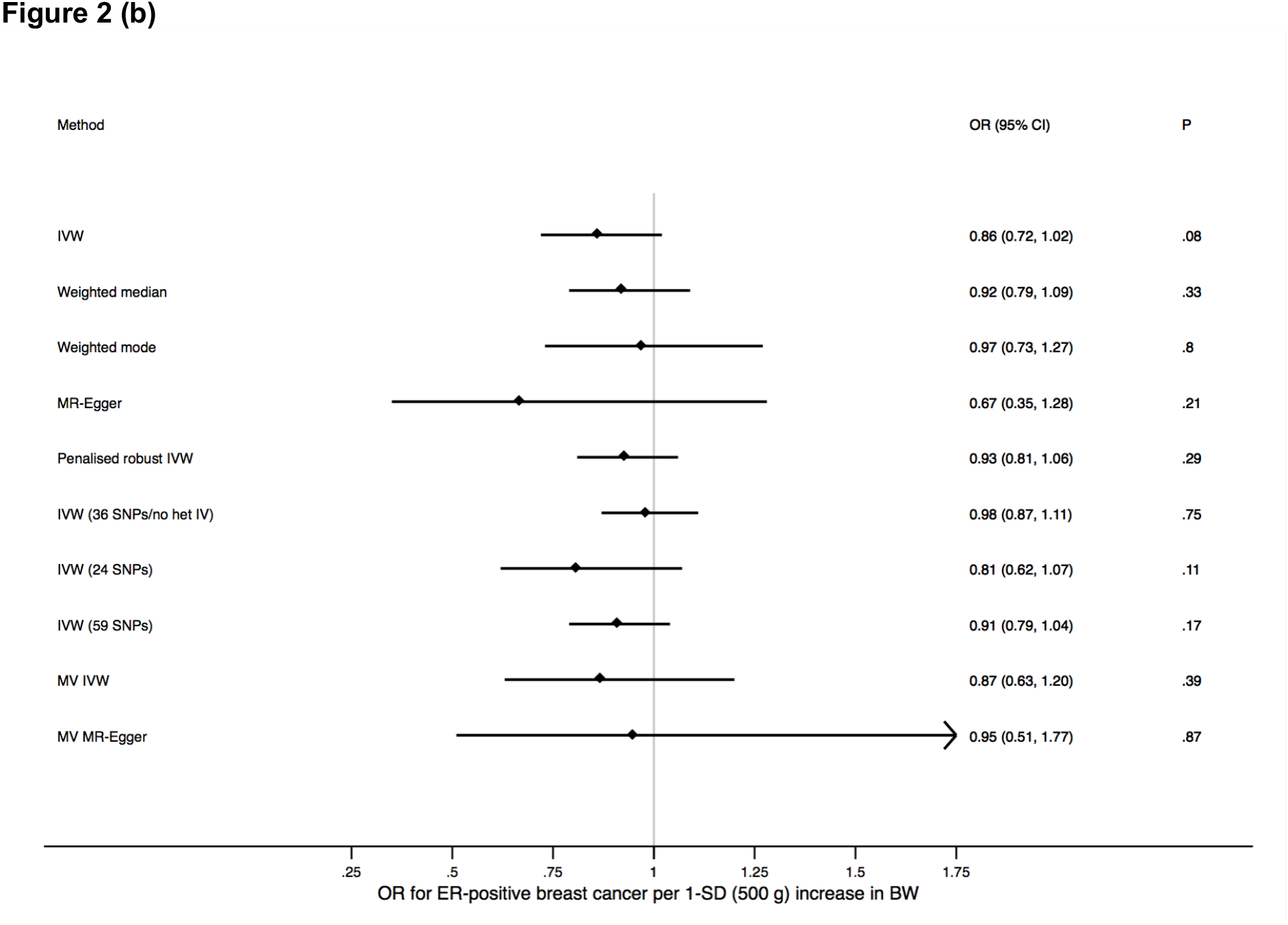
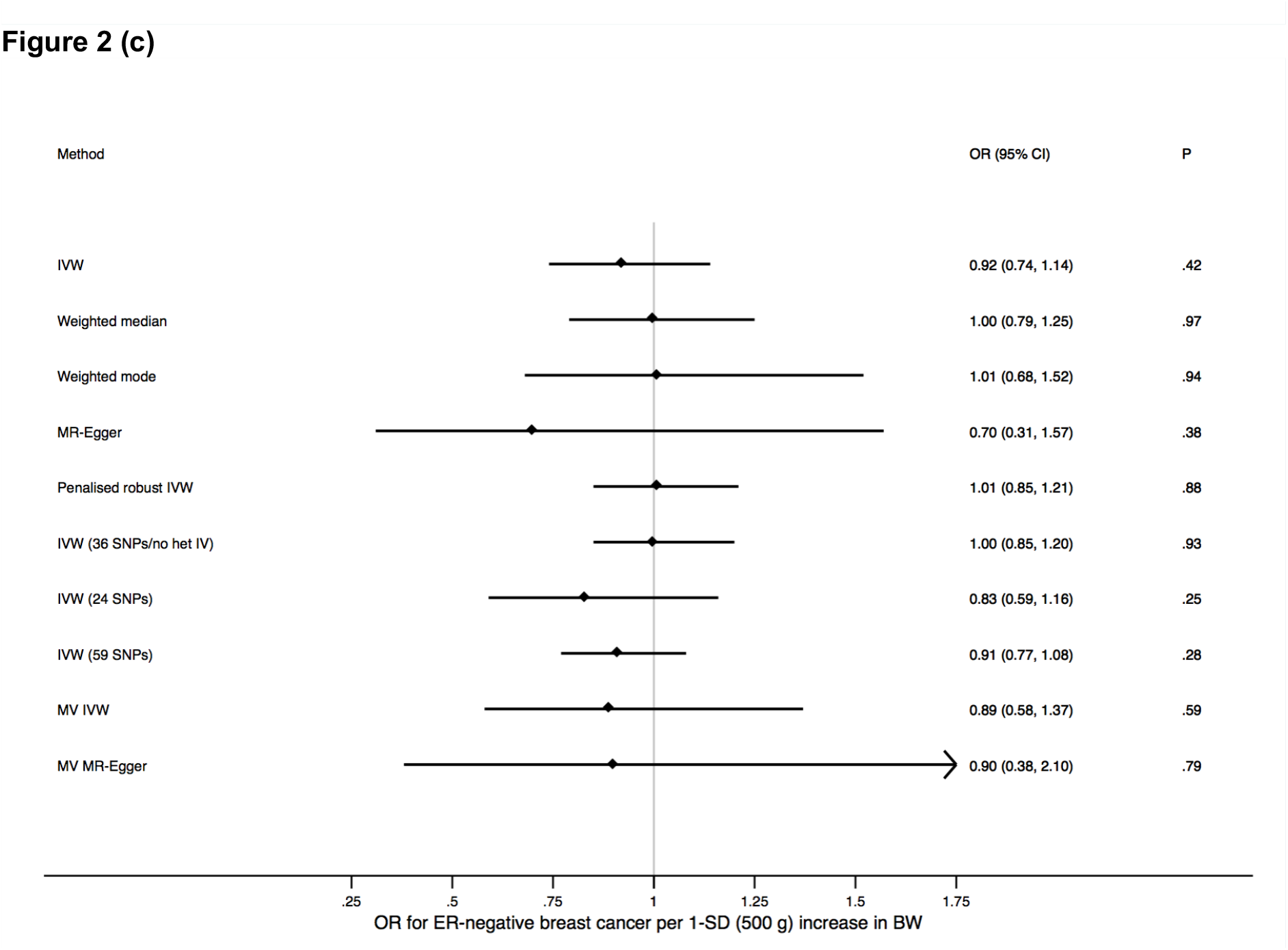
Forest plots of odds ratios (ORs) and 95% confidence intervals (CIs) for the association between birth weight (BW) and **(a)** overall breast cancer, **(b)** estrogen receptor (ER)-positive breast cancer, and **(c)** ER-negative breast cancer based on the different Mendelian randomisation approaches used in this study. Multivariable (MV) methods used the 49-SNP instrumental variable (IV) and all other methods, unless otherwise specified in the plots, used the 41-SNP IV. IVW indicates inverse-variance weighted regression, SD is standard deviation, and “no het” refers to no significant between-instrument heterogeneity at *P* < 0.05 based on Cochran’s Q.

Testing for heterogeneity between SNPs in the IV using Cochran’s Q indicated significant heterogeneity in Wald ratios for individual SNPs in the IV (*P* = 3.6 × 10^-16^ for overall breast cancer; *P* = 6.4 × 10^-11^ for ER-positive breast cancer; *P* = 6.5 × 10^-5^ for ER-negative breast cancer), suggesting that some of the 41 SNPs may exert disproportionately large effects on breast cancer risk some of which might act via pathways other than BW. Leave-one-out permutation (Supplementary Figure 1 (b)), Cook’s distance, and stepwise downward “model selection” consistently identified five SNPs with large effects on breast cancer that had an undue influence on the overall results (Supplementary Table 6). The stepwise downward approach suggested that an additional (sixth) SNP also contributed to the observed heterogeneity, while studentised residuals identified three of the five SNPs as outliers (Supplementary Table 6). Removing the five SNPs and repeating the standard IVW method using the remaining 36 SNPs that represented a more homogeneous genetic instrument for BW yielded an OR of 0.92 (95% CI: 0.83—1.02; *P* = 0.11; Cochran’s Q = 49.57 and its associated *P* = 0.052). Further reducing the potential impact of pleiotropy at the cost of losing power by using the 24-SNP IV constructed after removal of all SNPs associated at *P* < 9.2 × 10^-4^ (instead of *P* < 5 × 10^-8^) with at least one of seven putative breast cancer risk factors did not significantly change the primary IVW MR result (Figure 2 (a)). Conversely, using the full 59-SNP IV to leverage maximum statistical power did not meaningfully alter the primary result either (Figure 2 (a)). Results for each corresponding analysis specific to ER-positive and ER-negative breast cancer are presented in Figures 2 (b) and (c), Supplementary Figures 1 (c) to (f), and Supplementary Table 6. Across all analyses point estimates of the OR were either null or indicated an inverse relationship between BW and adult breast cancer, again contrary to the observational literature.

### Multivariable Mendelian randomisation

MR analysis of the 49-SNP IV for BW simultaneously adjusting for the genetically predicted effects of these SNPs on birth length, adult height, adult BMI, age at menarche, and age at menopause using the weighted regression-based framework provided estimates for the association between BW and overall breast cancer risk that were consistent with the univariable results (OR = 0.82; 95% CI: 0.60—1.11; *P* = 0.20; Figure 2 (a)). The estimates from the multivariable extension of MR-Egger were similar (Figure 2 (a)) and the corresponding MR-Egger intercept test was not significant (*P* = 0.89 for overall breast cancer; *P* = 0.75 for ER-positive breast cancer; *P* = 0.99 for ER-negative breast cancer). Results specific to ER-positive and ER-negative breast cancer are presented in Figures 2 (b) and (c).

## Discussion

We conducted a two-sample MR study using summary statistics from the largest genome-wide association meta-analyses data sets currently available for BW and for breast cancer susceptibility in adults. Our study provides no evidence to support the association between higher weight at birth and an increased risk of developing breast cancer in later life that was previously reported in observational studies. In fact, we found that higher genetically-predicted BW might, if anything, be associated with reduced breast cancer risk. Our results were robust to the application of different MR approaches, each with its own distinct set of assumptions.

A few observational studies suggest that the association between higher BW and increased future breast cancer risk is confined to premenopausal women but combined analyses of these studies show that this association does not differ by menopausal status [7,8]. ER-negative breast cancers are more common in premenopausal women and our MR study did not reveal any differences in the (lack of) association between genetically-predicted BW and breast cancer risk by ER-status (Figure 2 (b) and (c)). Some of the epidemiological literature also suggests that longer birth length may be a stronger risk factor for adult breast cancer than, and independently of, higher BW [7]. Birth length is highly correlated with BW and harder to measure as a phenotype making its measurement prone to error. Only two SNP-associations (rs905938 and rs724577) with birth length have been identified at genome-wide significance (*P* < 5 × 10^-8^) to date [31]. Both SNPs were also associated at *P* < 5 × 10^-8^ with BW and at *P* < 6 × 10^-8^ with adult height in the data sets that we used (Supplementary Table 3; the birth length-increasing alleles also increased BW and adult height). We did not use MR to assess the independent effect of birth length on adult breast cancer risk due to the availability of just two SNPs both of which are strongly pleiotropic and the consequent potential for unreliable estimates with such a genetic instrument.

While our point estimates for the effect of BW on breast cancer risk never reached statistical significance at the *P* < 0.05 level, they did uniformly indicate that the effect (if any) of higher BW on adult breast cancer might be protective (Figure 2). A possible protective effect of higher BW on adult breast cancer risk would be consistent with MR findings that genetically-predicted adult BMI is associated with reduction in breast cancer risk but, crucially, opposite in direction to estimates from combined analyses of observational studies investigating the BW-breast cancer link [7,8,54]. MR analysis has previously identified an association between higher BW and elevated BMI in adulthood and genome-wide genetic correlation analysis carried out as part of the Horikoshi et al. study also supported a positive correlation between BW and adult BMI [29,53]. Our MR results, however, reflect the effects of the genetic IV for BW on breast cancer independent of the effects of the IV on adult BMI since we removed adult BMI-associated SNPs from the IV before univariable analyses and adjusted for SNP associations with adult BMI as a covariate in multivariable analyses.

There are three aspects of the genome-wide association meta-analysis by Horikoshi et al. from which we obtained the BW lead SNPs to construct the IVs used in our MR study that are worth noting here [29]. First, our genetic IV with maximum power (59-SNP IV) explained only 2% of the variance of BW and therefore we were relatively underpowered to detect an odds ratio of 1.06 (or an even smaller effect) at the 5% significance level. Second, all autosomal SNPs genotyped on the array used by the UK Biobank (which was the largest sub-study in the genome-wide association meta-analysis by Horkioshi et al.) together explained approximately 15% of the variance of BW. Thus, it is likely that factors other than genetics account for the majority of the variance of BW. Third, the Horikoshi et al. study excluded individuals with extremes of BW (for the UK Biobank, which contributed nearly half the European sample, this was defined as < 2,500 g or > 4,000 g). Thus, the genetic IV we used may not adequately capture common genetic variants that only specifically affect the extremes of BW (i.e., only if such variants exist). This is relevant because in combined analyses of observational studies, statistically significant increases in adult breast cancer risk are only identified when comparing BW ≥ 4,000 g as an exposure to BW with 3,000 to 3,499 g or < 2,500 g as the reference categories [7,8]. However, it is highly unlikely that the genetic architecture of extreme BW completely differs from the genetic architecture of BW in general and therefore, our genetic IV likely predicts BW to some degree even at the extremes.

It has been suggested that insulin-like growth factor (IGF) signalling is one potential pathway linking increased BW to breast cancer risk in later life [3]. Haematopoietic stem or progenitor cells in neonatal cord blood have been used as a proxy for the size of the fetal mammary stem cell pool since the latter is impossible to measure. The level of haematopoietic stem cells in cord blood is positively correlated with IGF-1 levels in cord blood and with BW [5,55]. Cord blood IGF-1 concentrations have been found to be higher among Caucasian neonates than Chinese neonates and it has been hypothesised that these differences may be responsible, in part, for the differences in breast cancer risks observed between these two populations [56,57]. Lead SNPs near *IGF1* (rs7964361) and *IGF1R* (rs7402982) that encode IGF-1 and its receptor, respectively, were associated with birth weight at *P* < 5 × 10^-8^ and with adult height in women at *P* < 6 × 10^-4^ (Supplementary Tables 2 and 3). However, neither SNP was associated with overall or ER-positive or ER-negative breast cancer risks at *P* < 0.05 although the alleles that increased BW also conferred breast cancer risk (Supplementary Table 2; IVW MR analysis *P* = 0.59 for overall breast cancer risk when using just these two SNPs as an IV for BW). Thus, while there is an association between SNPs in the core IGF-1 signalling genes and BW, our data do not provide any evidence that higher BW as predicted by these two IGF pathway SNPs is associated with adult breast cancer risk. While data on the association between BW and breast cancer risk in non-European populations is scarce, it is also worth noting here that a small study from China failed to show any association [23].

Since genetically-predicted BW is not associated with increased adult breast cancer risk, a key unanswered question is whether the association between higher BW and increased breast cancer risk seen in observational studies is largely due to the BW-increasing effect of non-genetic factors such as maternal hormones and nutrition. In this regard, it is instructive to look at the results of an MR study that examined another well-known association between BW and a common non-communicable disease of adulthood. The association between low BW and coronary artery disease (CAD) risk in adults was first observed in 1989 (around the same time that the intrauterine origins of breast cancer were first proposed) and since then many observational studies have supported this association [58,59]. Underlying the BW-CAD association has been the hypothesis that stress induced by fetal malnutrition leads to intrauterine growth restriction, low BW, and metabolic reprogramming to maximise uptake of all available nutrition in later life culminating in a cardio-metabolic profile conducive for the development of CAD [60]. However, despite the contribution of the intrauterine environment to BW implicit in this chain of events, MR has identified an association between genetically-lower BW and increased adult CAD risk, in this case confirming the results from observational epidemiology [53]. Moreover, two MR analyses have also strongly supported an association between genetically-lower BW and increased adult type 2 diabetes risk [52,53]. Thus, while we are unable to completely rule out an association between the non-genetically-determined component of BW and breast cancer, the CAD and T2D findings indicate that genetically-predicted BW can help confirm observational associations with adult disease that may, at least partially, result from the non-genetic determinants of BW.

In conclusion, the association between higher BW and increased risk of developing breast cancer in adulthood has been identified in several observational studies published over the last three decades. However, revisiting this association using a comprehensive set of recently-developed MR methods and the largest available genomic data sets provides no evidence to suggest that the association is causal.

## Acknowledgements

This work builds on multiple, publicly available data sets. Data on birth weight and birth length were obtained from the Early Growth Genetics (EGG) Consortium. Data on adult height and adult body mass index were obtained from the Genetic Investigation of ANthropometric Traits (GIANT) Consortium. Data on age at menarche and age at menopause were obtained from the Reproductive Genetics (ReproGen) Consortium. Data on age at first birth and number of children ever born were obtained from the Social Science Genetic Association Consortium (SSGAC). Data on type 2 diabetes were obtained from the DIAbetes Genetics Replication And Meta-analysis (DIAGRAM) Consortium. Data on breast cancer were obtained from the Breast Cancer Association Consortium (BCAC).

